# PERK arm of UPR selectively regulates ferroptosis in colon cancer cells by modulating the expression of SLC7A11 (System Xc-)

**DOI:** 10.1101/2023.03.28.534659

**Authors:** Krishan Kumar Saini, Priyank Chaturvedi, Abhipsa Sinha, Manish Pratap Singh, Muqtada Ali Khan, Ayushi Verma, Mushtaq Ahmad Nengroo, Saumya Ranjan Satrusal, Sanjeev Meena, Akhilesh Singh, Sameer Srivastava, Jayanta Sarkar, Dipak Datta

**Author notes:** Corresponding Author: Dipak Datta, Division of Cancer Biology, CSIR-CDRI, B.S. 10/1, Sector 10, Jankipuram Extension, Sitapur Road, Lucknow-226031, India, (Extn-4347/48).

## Abstract

Ferroptosis, a genetically and biochemically distinct form of programmed cell death, is characterised by an iron-dependent accumulation of lipid peroxides. Therapy-resistant tumor cells display vulnerability toward ferroptosis. Endoplasmic Reticulum (ER) stress and Unfolded Protein Response (UPR) play a critical role in cancer cells to become therapy resistant. Tweaking the balance of UPR to make cancer cells susceptible to ferroptotic cell death could be an attractive therapeutic strategy. To decipher the emerging contribution of ER-stress in the ferroptotic process, we observe that ferroptosis inducer RSL3 promotes UPR (PERK, ATF6, and IRE1α), along with overexpression of cystine-glutamate transporter SLC7A11 (System Xc^-^). Exploring the role of a particular UPR arm in modulating SLC7A11 expression and subsequent ferroptosis, we notice that PERK is selectively critical in inducing ferroptosis in colorectal carcinoma. PERK inhibition reduces ATF4 expression and recruitment to the promoter of *SLC7A11* and results in its downregulation. Loss of PERK function not only primes cancer cells for increased lipid peroxidation but also limits in vivo colorectal tumor growth, demonstrating active signs of ferroptotic cell death *in situ*. Further, by performing TCGA data mining and using colorectal cancer patient samples, we demonstrate that the expression of *PERK* and *SLC7A11* is positively correlated. Overall, our experimental data indicate that PERK is a negative regulator of ferroptosis and loss of PERK function sensitizes colorectal cancer cells to ferroptosis. Therefore, small molecule PERK inhibitors hold huge promise as novel therapeutics and their potential can be harnessed against the apoptosis-resistant condition.

## Introduction

Cancer is a leading cause of death worldwide; in 2020, there were 19.3 million new cases of all types of cancer in which more than half of the patients died. Colorectal cancer is the third most commonly diagnosed cancer (10.0%) and the second leading cause of death (9.4%) worldwide in both males and females [1]. Therapy resistance is the key to tumor relapse and subsequent tumor-associated mortality. Evasion of apoptosis is one of the important hallmarks of cancer cells and mechanisms behind the same have enormous therapeutic potential in the context of current cancer research [2]. Recently, we have shown that how intracellular CXCR4 protein and epigenetic modulator EZH2 promote therapy resistance, CSC properties and metastasis in colorectal and breast cancer [3–5]. Recent reports also suggest that these resistant cancer cells are vulnerable to iron-mediated cell death or ‘Ferroptosis’ [6,7].

As originally discovered by the Stockwell group, ferroptosis is morphologically, biochemically and genetically distinct from apoptosis, necroptosis, and autophagy and depends on intracellular iron [8,9]. Selenoprotein Glutathione peroxidase 4 (GPx4), cystine/glutamate antiporter (System Xc⁻) and enzyme Acyl-CoA synthetase long-chain family member 4 (ACSL4) are known to be the key modulators of ferroptotic process [10–12]. GPx4 is the critical enzyme that can reduce lipid hydroperoxides within biological membranes; hence ferroptosis can be induced by the treatment of small molecule GPx4 inhibitor RSL3 (Ras Selective Lethal) treatment [13,14]. System Xc^-^ imports cystine in exchange for glutamate; cystine further takes part in the biosynthesis of glutathione (GSH) [15]. ACSL4 modulates ferroptosis sensitivity by shaping the cellular lipid composition of the cell [16].

Though therapy-resistant cells have vulnerability to ferroptotic cell death, these cells are proficient in handling therapeutic insults by adapting to cellular stress caused by Unfolded Protein Response (UPR)[17]. UPR is initiated by three transmembrane proteins PERK (Protein kinase RNA-like endoplasmic reticulum kinase), ATF6 (activating transcription factor 6), IRE1α (inositol-requiring enzyme 1α), which are commonly known as three arms of UPR [18,19]. These three proteins are responsible for maintaining cell survival and homeostasis by regulating protein folding in response to UPR [20]. These proteins have an ER-luminal domain believed to sense the protein misfolding; alteration in this domain in response to stress also changes the oligomerization state of these UPR proteins. PERK and IRE1α only need the oligomerization of the luminal domain to be activated; an ER chaperone, Bip/GRP78 binding, keeps these two proteins inactive by maintaining these proteins in ER membrane in unstressed cell conditions [21,22]. PERK has a single kinase domain that limits eIF2α activity by phosphorylating it, resulting in global translation attenuation, whereas ATF4 translation is selectively upregulated when active eIF2α is limiting [23]. IRE1α possesses kinase and RNase activity; when activated, IRE1α cleaves *XBP1*, and *XBP1* then activates transcription of many ER-related genes that regulate protein folding in the ER [24,25]. ATF6 is translocated to the Golgi lumen and cleaved by site-1 protease (S1P) and Site-2 protease (S2P) to produce ATF6 (N). ATF6 (N) and XBP1 promote protein transcription, which increases ER size and protein folding capacity [26,27]. These transcriptional processes work in concert as homeostatic feedback loops to reduce ER-stress. If the amount of misfolded protein is declined, UPR signaling is decreased and the cell survives.

The influence of UPR in regulating tumor cell apoptosis and autophagic cell death is vastly studied in the literature [28,29]. However, very little is known regarding the role of UPR in modulating ferroptosis. Here, we reveal that ferroptosis inducer RSL3 promotes an enormous amount of ER-stress in colorectal cancer cells. In the course of dissecting the impact of three independent UPR arms in modulating RSL3 induced ferroptosis, we observe that among all three arms of UPR, PERK selectively prevents ferroptotic tumor cell death. Further analysis suggests that PERK activation and downstream signaling protects tumor cells from ferroptosis through SLC7A11 upregulation. Similarly, loss of PERK function results in reduced colorectal tumor growth in vivo with increased ferroptosis. Finally, TCGA data mining and expression analysis of colorectal cancer patient-derived tumors with their matched normal counterpart display a positive correlation between *PERK* and *SLC7A11* expression signifying the clinical relevance of our finding.

## Results

### PERK arm of UPR positively regulates SLC7A11 (System Xc⁻) expression in colorectal cancer cells

The relationship between apoptosis/autophagic cell death and UPR or ER-stress is well established. However, the contribution of UPR in modulating ferroptotic cell death in colorectal cancer remains elusive. To investigate the above relationship, first, we tested the impact of ferroptosis inducer (RSL3) on three different arms of UPR (PERK, ATF6 and IRE1α) in two different colorectal cancer cells (HT29 and SW620). As shown in Figures 1A and 1B, classical ferroptosis inducer RSL3 promotes the expression of all three arms of UPR in both the cells. We also find similar upregulation of downstream effector proteins of UPR such as ATF4, Bip/GRP78 and XBP1 (Supplementary Figure 1) in HT29 and SW620 cells. Further, we evaluated its effect on the expression of ferroptosis signature genes, such as *GPx4, ACSL4*, and *SLC7A11*. As observed in Figure 1C, RSL3 treatment inhibits GPx4 expression as expected, being a GPx4 inhibitor, whereas the expression of SLC7A11 is markedly upregulated in HT29 cells, having no impact on ACSL4 expression. Similar results are observed in another colon cancer cancer (SW620) cells (Supplementary Figure 2). Further, we sought to determine the role of three different UPR arms in the expression of ferroptosis signature genes *GPx4, ACSL4*, and *SLC7A11* in HT29 colorectal cancer cells. In Figure 1D-1F, we observe individual knockdowns of each UPR arm like PERK, ATF6 and IRE1α in HT29 cells, resulting in non-noticeable changes in protein expression of ferroptotic genes except PERK knockdown markedly reduces the expression of SLC7A11 protein as compared to control. Since RSL3 was shown to upregulate SLC7A11 expression at basal condition, we evaluated the effect of RSL3 on control and PERK KD HT29 cells. As shown in Figure 1G, PERK knockdown strongly prevented RSL3 induced SLC7A11 overexpression compared to the control. Similarly, compared to vehicle control, PERK inhibitor alone (Figure 1H) reduces SLC7A11 expression and also mitigates RSL3 induced SLC7A11 expression (Figure 1I). Therefore, either the loss of PERK protein or functional loss of PERK kinase activity through PERK inhibitor dampens SLC7A11 expression not only at basal condition but also abrogates RSL3 mediated SLC7A11 induction. Additionally, we cultured EV and PERK knockdown HT29 cells for 72 hours in cystine^+^ and cystine^-^ conditions and observed that SLC7A11 expression is upregulated in cystine^-^ conditions compared to cystine^+^ control. The above phenomenon is significantly abolished in PERK knockdown cells (Figure 1J), suggesting that PERK is indispensable for SLC7A11 upregulation in cystine-starved conditions. Further, as shown in the Figure 1K-1L, pictorial representation and densitometric quantification show a significant reduction in cell number in PERK knockdown condition as compared to respective control when cells are cultured in the absence of cystine (Figure 1K-1L). Therefore, either the loss of PERK protein or functional loss of PERK kinase activity through PERK inhibitor dampens SLC7A11 expression not only at basal condition but also abrogates RSL3 mediated SLC7A11 induction.

**Figure 1:**
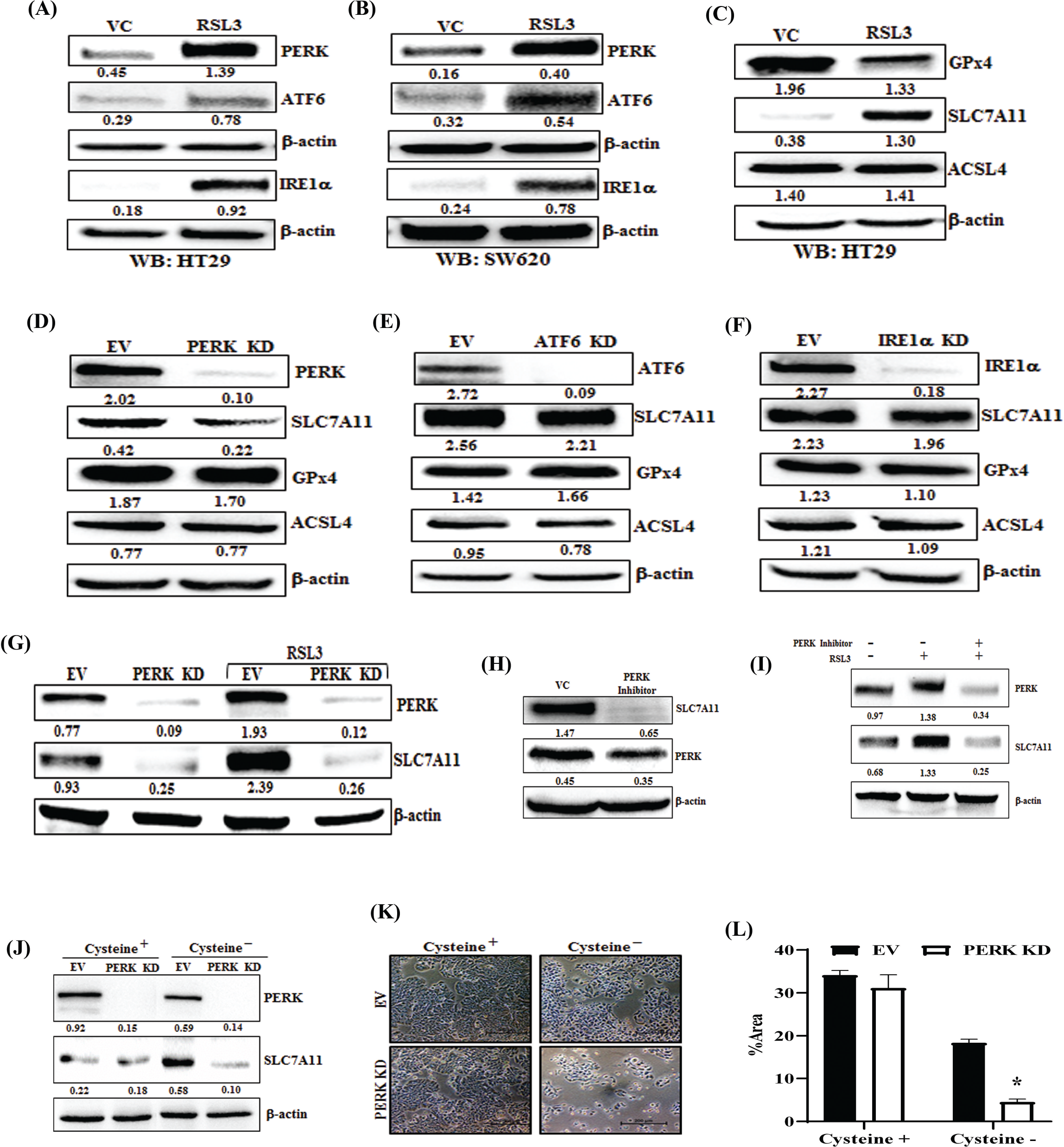
Ferroptosis inducer RSL3 causes UPR and the PERK arm of UPR regulates SLC7A11 expression. (A) HT29 and (B) SW620 cells were either treated with 1μM RSL3 or vehicle control (VC) for 24 hours and protein lysates were prepared for western blot analysis. Immunoblot shows the expression for classical UPR marker proteins, i.e., PERK, ATF6, and IRE1α. (C) Major ferroptosis regulator proteins GPx4, SLC7A11, and ACSL4 in HT29 cells. (D-F) Immunoblot analysis of PERK, ATF6, IRE1α, SLC7A11, GPx4, and ACSL4 in (D) PERK knockdown (KD), (E) ATF6 KD and (F) IRE1α KD in HT29 cells with respective empty vector (EV). (G) Western blot shows the expression of PERK and SLC7A11 in EV and PERK KD HT29 cells that were either treated with vehicle or 1μM RSL3 for 6 hours. (H-I) Immunoblot analysis of PERK and SLC7A11 in (H) VC and PERK inhibitor (GSK2656157) treated and (I) VC, PERK inhibitor and RSL3 treated HT29 cells. (J-L) EV and PERK KD HT29 cells were cultured in RPMI 1640 in the presence of cystine (cystine+) or absence (cystine-) conditions for 72 hours and subjected to Western blot analysis for the expression of (J) PERK and SLC7A11 and represented as (K) photomicrographs and (L) subsequent quantitative analysis of images are shown. β-actin was used as a loading control in all immunoblot studies. Densitometric quantifications are shown under each respective blot of all Western blot images.

### PERK loss of function sensitizes colorectal cancer cells to ferroptosis

SLC7A11 (System Xc⁻) is a vital membrane transporter that imports cystine into the cytosol in exchange for glutamate, while glutathione synthetase and γ-glutamylcysteine synthetase synthesize the antioxidant and GPx4 substrate glutathione. Our prior findings show that loss of PERK reduces System Xc⁻ expression, implying that this down-regulation might play a role in ferroptosis regulation. So, we treated different UPR arm knockdown HT29 cells with RSL3 and observed that PERK knockdown cells are more selectively sensitive to RSL3 mediated cell death than vehicle control, ATF6 KD, or IRE1α KD cells (Figure 2A). PERK knockdown in SW620 cells produces similar results (Figure 2B). We treated HT29 cells with erastin, another potent ferroptosis inducer, and observed sensitization in PERK knockdown cells as compared to control (Supplementary Figure 3). To further confirm the above observations, we seeded equal numbers of untagged HT29 EV (white) and chilli-luc tagged PERK (red) knockdown cells and cultured in the absence and presence of RSL3 for 3 days. FACS analysis of Day 0 and Day 3 cells shows a marked reduction of chilli luc tagged PERK (red) knockdown cells compared to untagged white control cells, again suggesting that loss of PERK sensitizes cytotoxic function of RSL3 (Figure 2C). However, we did not observe any sensitisation of PERK KD cells against classical chemotherapeutic drugs like paclitaxel, 5-fluorouracil and oxaliplatin (Supplementary Figure 4). To ascertain the above findings, we treated HT29 cells with classical FDA-approved drugs such as paclitaxel, 5-fluorouracil, doxorubicin, and oxaliplatin and observed no significant difference in cytotoxic potential of these drugs in control and PERK knockdown cells (Figure 2D). Another colon cancer cell line SW620 also showed similar results for the above drugs in both conditions (Supplementary Figure 5). The above findings suggest that sensitization observed by PERK knockdown is specific to RSL3. Now, we sought to determine whether RSL3 mediated sensitization of cell death in the absence of PERK is due to the induction of apoptosis or ferroptosis. To find out the same, we treated PERK knockdown cells with RSL3 in the presence or absence of apoptosis inhibitor (pan-caspase inhibitor Z-VAD-FMK) or ferroptosis inhibitor (ferrostatin-1) and observed that only ferroptosis inhibitor (ferrostatin-1) significantly rescues cytotoxic impact of RSL3 (Figure 2E). The above findings demonstrate the role of ferroptosis in PERK knockdown-mediated sensitization of cell death. Additionally, there was no significant difference in cell growth inhibition with RSL3 and pan-caspase inhibitor, so classical apoptotic cell death has a minimal role associated with RSL3-induced cell death. Ferroptosis is distinct from other types of programmed cell death and is characterised by the accumulation of lipid peroxides that can be detected by BODIPY-C11 staining of cells [30]. To check the level of lipid peroxidation after RSL3 treatment, we treated HT29 (Figure 2F, left and right panel) and SW620 (Figure 2G, left and right panel) EV and PERK knockdown cells with RSL3 for 24 hours and subjected them to BODIPY-C11 staining (details described in materials and methods), we observe a marked increase in excitation shift of BODIPY-C11 staining in PERK knockdown cells as compared to control, suggesting that loss of PERK promotes lipid peroxidation and ferroptosis in colorectal cancer cells. 4-HNE and MDA are known lethal by-products of lipid peroxidation or ferroptosis, formed during the enzymatic and non-enzymatic breakdown of AA (arachidonic acid) and other PUFAs (polyunsaturated fatty acids) [31,32]. Here, we assessed the expression of 4-HNE and MDA in control, PERK KD and PERK inhibitor (GSK2656157) treated cells and found robust upregulation of both in PERK KD (Figure 2H) and PERK inhibitor (Figure 2I) treated cells as compared to respective controls. The above observations highlight the importance of PERK as a negative regulator of ferroptosis in colorectal cancer and also suggest that apoptotic cell death has a minor role in PERK mediated ferroptosis modulation.

**Figure 2:**
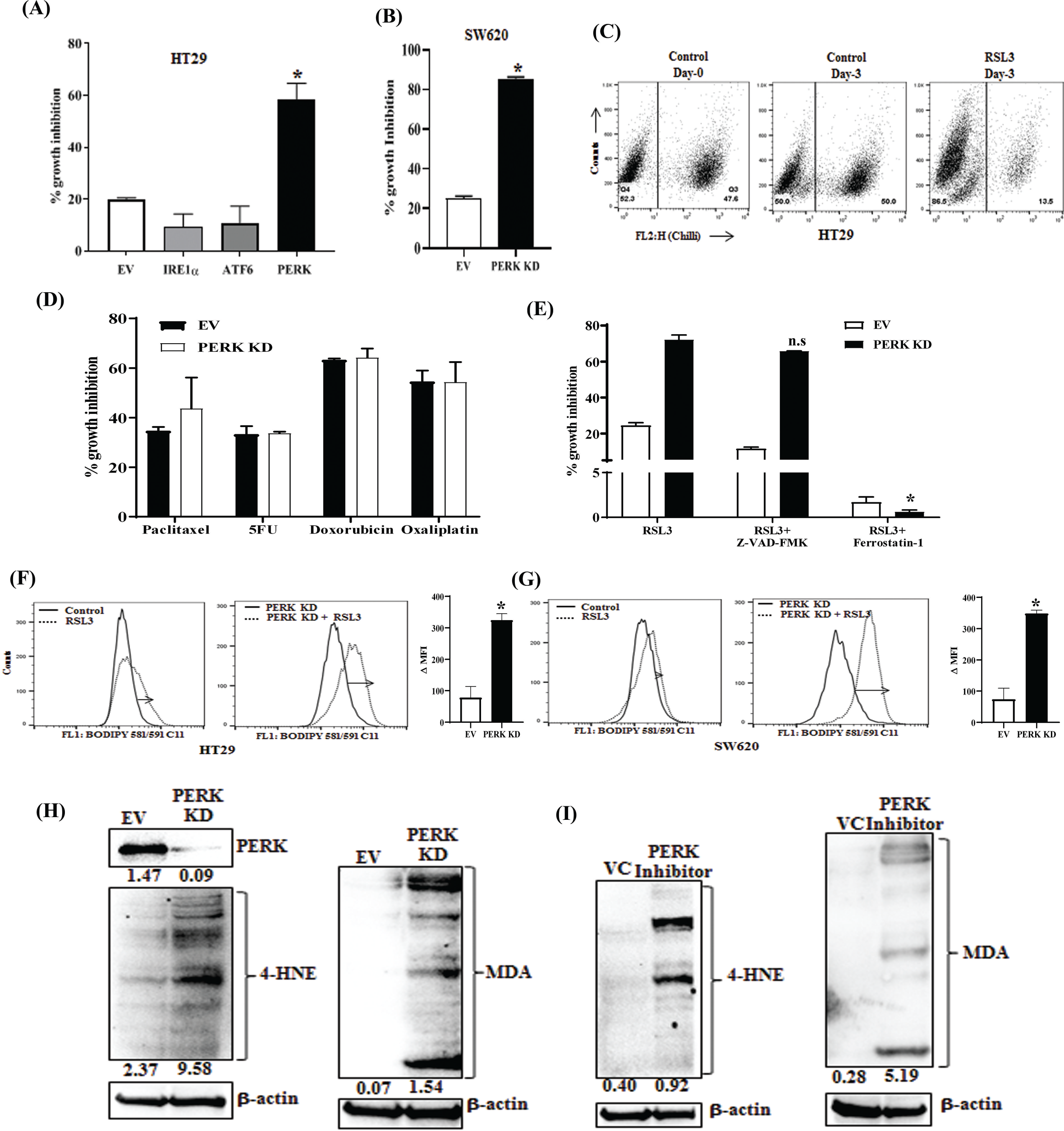
Loss of PERK function promotes ferroptosis in colorectal cancer. (A) EV and PERK, ATF6, and IRE1α KD HT29 cells were treated with either vehicle or 1μM RSL3 for 48 hours and subjected to SRB assay to evaluate the cytotoxic effect of the same. (B) EV and PERK knockdown SW620 cells were treated with vehicle or 1μM RSL3 for 48 hours and the SRB assay was performed. For A and B, percent growth inhibition was tabulated. *p < 0.05; compared to EV. (C) HT29 EV (untagged) and chilli-tagged PERK KD (red) cells were mixed in equal numbers and subjected to flow cytometric analysis on day 0 and day 3 following treatment with vehicle control or 500nM RSL3. Analysed cell populations are shown in the FACS plot. (D) Control and PERK KD HT29 cells were treated with either vehicle control or with different chemotherapeutic drugs paclitaxel (20nM), 5-Fluorouracil (100μM), Doxorubicin (10μM), Oxaliplatin (50μM) for 48 hours and cytotoxic impact of these drugs was evaluated via SRB assay. Percent growth inhibition was tabulated. (E) HT29 EV and PERK KD cells were treated with either 1μM RSL3 alone or in combination with (25μM) Z-VAD-FMK (pan-caspase inhibitor) or (10μM) ferrostatin-1 (ferroptosis inhibitor) and percent growth inhibition in different groups was estimated by SRB assay. In A, B, D and E Columns represent an average of triplicate readings of samples; error bars ± S.D. (F-G left panel) Control (EV) and PERK KD of (F) HT29 and (G) SW620 cells were treated with 1μM RSL3 or vehicle control for 24 hours followed by BODIPY C11 staining (Lipid peroxidation sensor) and cells were analysed by FACS (detailed description materials and methods section). Histogram overlays show BODIPY C11 positivity correlating with lipid peroxidation levels in respective groups. (F-G) Right Panels, respective delta mean fluorescence intensity (MFI) of the cells, stained for BODIPY C11. The delta mean was calculated by subtracting the mean fluorescence intensity of the control from that of the RSL3 treated cells. Columns represent an average of duplicate readings of samples; error bars ± S.D. *p < 0.05; compared to EV (control). (H-I) Immunoblots representing 4-Hydroxynonenal (4-HNE) and malondialdehyde (MDA) conjugated protein expression in (H) HT29 EV and PERK KD cells and (I) HT29 VC and PERK inhibitor (GSK2656157) treated (24hours) cells. β-actin was used as a loading control. Western blot densitometric quantifications are shown below the respective blots.

### PERK regulates the expression and recruitment of transcription factor ATF4 to the promoter of SLC7A11 in the course of ferroptosis modulation

PERK mediated robust upregulation of SLC7A11 protein expression prompted us to evaluate the impact of PERK in *SLC7A11* transcriptional regulation. First, we observed that PERK knockdown severely reduces mRNA expression of *SLC7A11* in colorectal cancer cells, as observed in Figure 3A. As ATF4 is a critical downstream transcription factor of *PERK*, we further determined the level of ATF4 protein in control and PERK knockdown cells and found that level of ATF4, as well as SLC7A11, are markedly downregulated in PERK knockdown cells as compared to control (Figure 3B). Moreover, to understand the direct regulation of ATF4 on SLC7A11 expression, we made stable knockdown of ATF4 in HT29 cells and observed that SLC7A11 protein expression is robustly downregulated in ATF4 knockdown cells as compared to control (Figure 3C). Further, we wanted to evaluate the contribution of ATF4 recruitment in PERK mediated downregulation of SLC7A11 expression. As shown in Figure 3D, the *SLC7A11* promoter has three putative ATF4 binding sites predicted by the publicly available software Eukaryotic Promoter Database (https://epd.epfl.ch/). In our ChIP assay (Figure 3E), we observed marked selective enrichment of ATF4 at the −0.3kb site upstream of the *SLC7A11* promoter in control cells, which was found to be significantly reduced following PERK inhibitor treatment. Loss of ATF4 recruitment at the *SLC7A11* promoter following PERK inhibition prompted us to check its role in modulating *SLC7A11* transcription. In our dual luciferase assay, we transfected the SLC7A11 promoter luciferase construct in HT29 cells (-0.6 kb from the TSS, cloned in the luciferase assay reporter vector pGL4.12), indicating that PERK inhibitor reduces SLC7A11 transcription (relative luciferase activity) in a dose-dependent manner (Figure 3E). The above data suggest that ATF4 functions as a transcription factor in PERK mediated regulation of *SLC7A11* expression. To assess the phenotypic impact of the PERK-ATF4-SLC7A11 axis, we treated control and ATF4 KD cells with RSL3 and performed cell viability assay and BODIPY C11 staining. As observed in Figure 3F, ATF4 loss not only makes cells prone to death in response to RSL3 but also promotes increased ferroptosis compared to control (Figure 3G). Altogether, our experimental results clearly indicate that PERK regulates ferroptosis through *SLC7A11* expression by modulating the recruitment of transcription factor ATF4 to its promoter.

**Figure 3:**
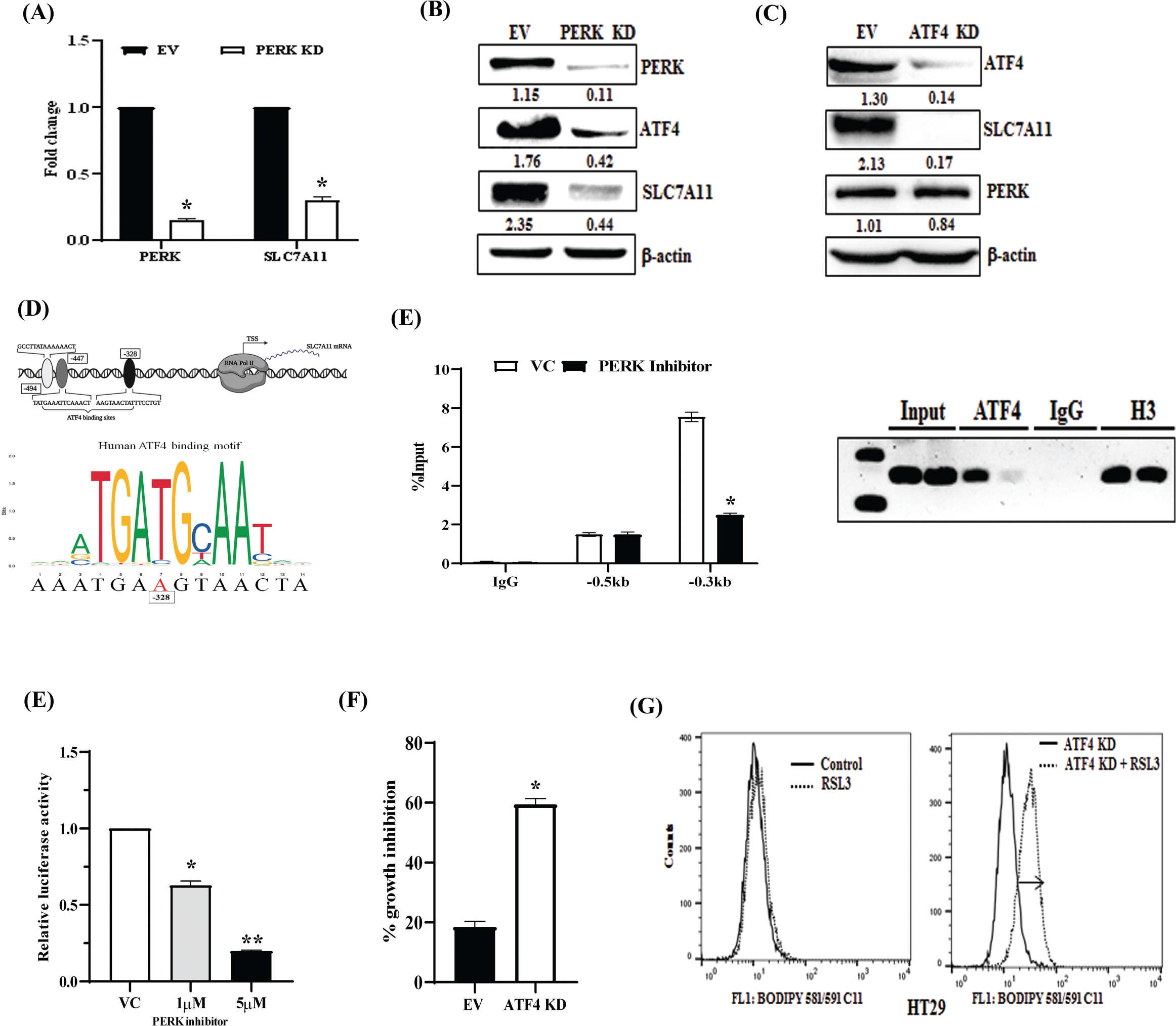
PERK-ATF4 axis regulates *SLC7A11* expression and colorectal cancer ferroptosis. (A) Total RNA was isolated from HT29 EV and PERK KD stable cells, reverse transcribed, and subjected to qRT-PCR analysis for *PERK* and *SLC7A11* mRNA expression. Fold change in mRNA expression is represented in the bar graph. Data is representative of three independent experiments, resulting from different samples; Columns, the average value of mRNA expression of *PERK* and *SLC7A11*, respectively; bars ± SD. *, p < 0.05, compared with EV. (B-C) Immunoblots show the expression of PERK, ATF4, and SLC7A11 in (B) HT29 EV and PERK KD or (C) HT29 EV and ATF4 KD cells. β-actin was used as the loading control. Western blot densitometric quantifications are displayed underneath each blot. (D) Diagrammatic representation of *SLC7A11* promotor showing (Top) putative ATF4 DNA binding sites and transcription start site (TSS) with RNA pol II, (Down) Human ATF4 binding motif of *SLC7A11* promoter on predicted binding site (-0.3kb upstream from TSS) that is publicly available at JASPAR database (http://www.jaspar.genereg.net). (E) ChIP (Details described in the Methods section) was performed in vehicle control, and PERK inhibitor treated HT29 cells using anti-ATF4 and IgG antibodies and then examined by real-time qPCR using primer pairs targeting predicted −0.3 kb and −0.5kb sites upstream from TSS of the *SLC7A11* gene. Fold change enrichment for ATF4 with respect to % input was shown; Data is representative of three independent experiments resulting from different samples; Columns, the average value of percentage enrichment compared to input; bars ± SD. *, p < 0.05, compared with vehicle control. Photomicrograph of Gel showing conventional PCR validation of ChIP experiments. Lanes are vehicle control and treatment, respectively, for each group of the ChIP sample. (G) The HEK293 cells were transfected with −0.3kb upstream of *SLC7A11* promoter luciferase construct plasmid followed by treatment with vehicle or 1μM and 5μM of PERK inhibitor for 24 hours and cells were harvested for luciferase activity (detailed description in Methods Section). Columns, the average value of relative firefly luciferase activity compared to Renila luciferase activity derived from triplicate readings of different samples; bars ± SD. **, p < 0.01, compared with vehicle control. (H) ATF4 KD and EV HT29 cells were treated with 1μM RSL3 for 48 hours, and SRB assay was performed. Percent growth inhibition was tabulated, Columns, an average of triplicate readings of samples; error bars ± S.D. *p < 0.05; compared to EV. (I) HT29 EV and ATF4 KD cells were treated with 1μM RSL3 for 6 hours and analysed by flow cytometry after staining with BODIPY C11. Histogram overlays show lipid peroxidation levels in respective treatment groups. Each data is representative of three independent experiments.

### Loss of PERK inhibits tumor growth and demonstrates active signs of ferroptotic tumor cell death in vivo

Considering the above-mentioned findings, we sought to look into the influence of PERK knockdown on tumor progression in vivo. We inoculated EV (control) and PERK KD HT29 cells (2×10^6^ respectively) subcutaneously in nude mice and monitored the tumor progression twice a week for up to 6 weeks. As observed in Figure 4A-4C, compared to the control, PERK KD resulted in a marked reduction of tumor volume (Figure 4A, 4B) and weight (Figure 4C). Further, we performed western blot analysis of the harvested tumors to investigate the effect of PERK loss of function on the expression of SLC7A11 as well as hallmark ferroptotic markers such as 4-HNE and MDA in vivo. As observed in Figure 4D, PERK knockdown is maintained in vivo tumors and loss of PERK strongly reduces SLC7A11 expression (Left Panel) but promotes marked expression of both ferroptotic markers such as 4-HNE (Middle Panel) and MDA (Right Panel). To further validate the above finding, we performed immunohistochemistry (IHC) staining using MDA and 4-HNE antibodies in control and PERK knockdown tumors. We observed that compared to the control, PERK KD tumors display robust overexpression of 4-HNE and MDA (Figure 4E). Further, we have calculated the IHC ATM (Averaged Threshold Measure) score by measuring the DAB intensity in respective IHC images, as represented in Figure 4F-G, where we find that PERK KD tumors have robustly high ATM scores for both 4-HNE and MDA staining as compared to the control tumor. Together, our data indicate that PERK has an immense impact on tumor growth and genetic inhibition of PERK results in significant loss of tumor growth, displaying the elevated levels of 4-HNE and MDA proteins suggest active ferroptotic tumor cell death in the course of PERK mediated tumor growth reduction.

**Figure 4:**
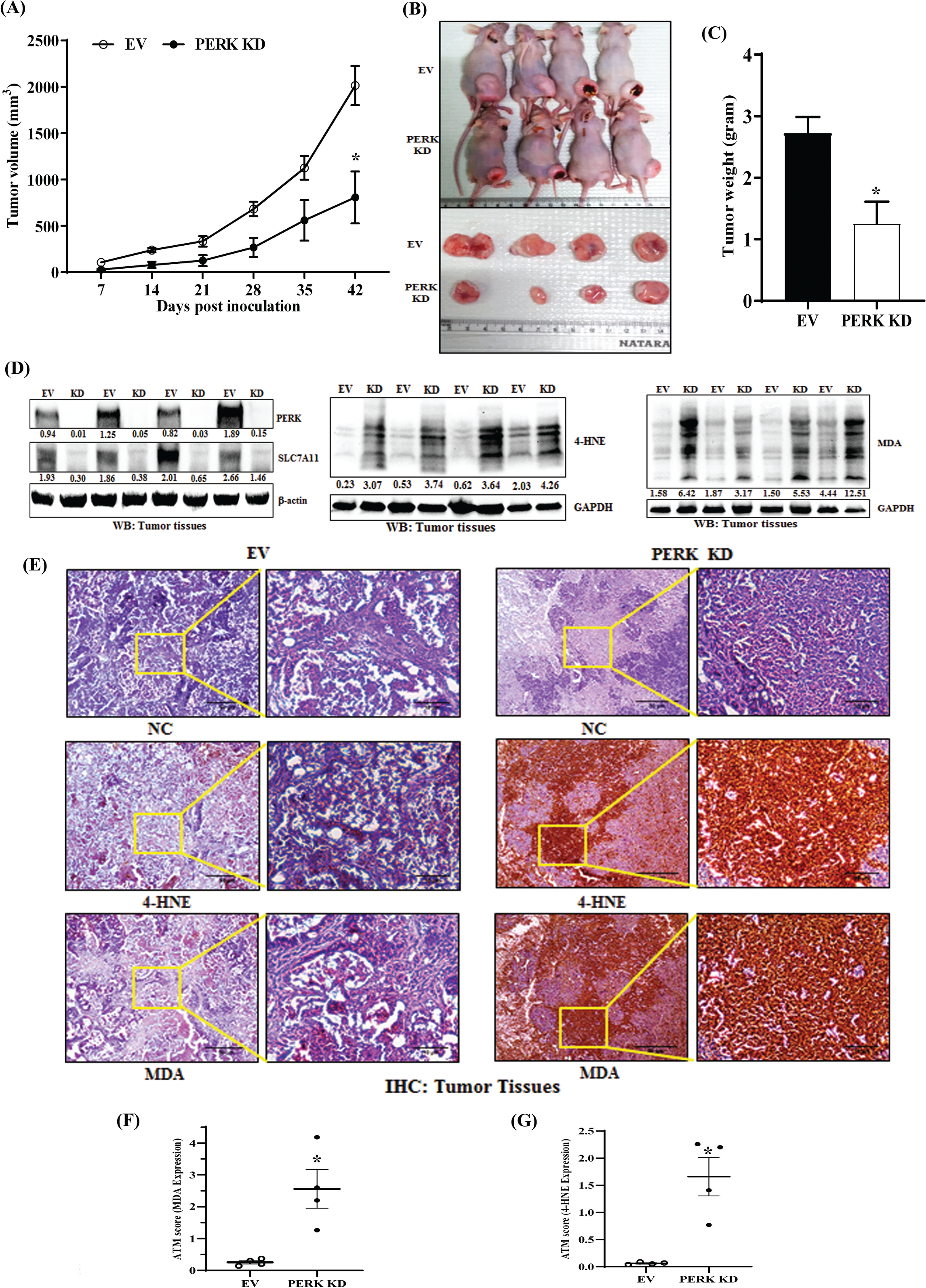
Loss of PERK has compromised in vivo colorectal tumor growth due to increased ferroptosis. 2 x 10^6^ HT29 EV and PERK KD cells in 100μl PBS were injected subcutaneously in the flanks of the right hind leg of 4–6 weeks old Crl: CD1-Foxn1nu mice in two different groups for each condition. Tumor volumes were measured twice a week with a caliper. (A) Tumor progression of the same is shown in the graph. Each point indicates the average tumor volumes at a particular time; error bars ± SEM (n = 4 for each group); *p < 0.05 compared to control tumors. (B) Photographs of tumor-bearing mice (top) and harvested tumors (bottom) from respective groups were shown. (C) The average tumor weight of each group is shown in the graph. error bars ± SEM (n = 4 for each group); *p < 0.05 compared to control tumors. (D) Harvested tumors were lysed and subjected to western blot analysis to visualize the protein expression of PERK and SLC7A11, MDA (malondialdehyde) and 4-HNE (4-Hydroxynonenal). β-actin and GAPDH were used as loading control. Western blot densitometric quantifications are shown below the respective blots. (E) Immunohistochemistry was conducted to detect MDA and 4-HNE in formalin-fixed paraffin-embedded serial sections of harvested tumors with respective antibodies. Representative photomicrographs are shown at 10X and 40X magnifications. Scale bar, 50μm (10X) or 10μm (40X). (F-G) Quantitative ATM scores for the expression of (F) MDA and (G) 4-HNE are represented as scatter plots; Error bar, +/− SEM, *p-value, <0.05, compared to expression in control tumors.

### Expression of *PERK* and *SLC7A11* are positively correlated in human colorectal cancer

To draw a clinical correlation between *PERK* and *SLC7A11* expression, we performed TCGA data mining using the UCSC Xena browser (https://xenabrowser.net/). We accessed mRNA expression data of 471 patients in the GDC TCGA COAD cohort of the database and drew a correlation plot between *PERK* and *SLC7A11* expression, and calculated Pearson’s correlation (R), where we found a positive correlation between *PERK* and *SLC7A11* expression (Figure 5A). Further, we confirmed a positive correlation between *PERK* and *SLC7A11* expression in 15 human colorectal cancer patients by performing Real-Time-PCR analysis of CRC tumors and their respective matched normal counterparts (Figure 5B). As shown in Figure 5C-5D, we find that both PERK and SLC7A11 are overexpressed in human CRCs compared to normal counterparts.

**Figure 5:**
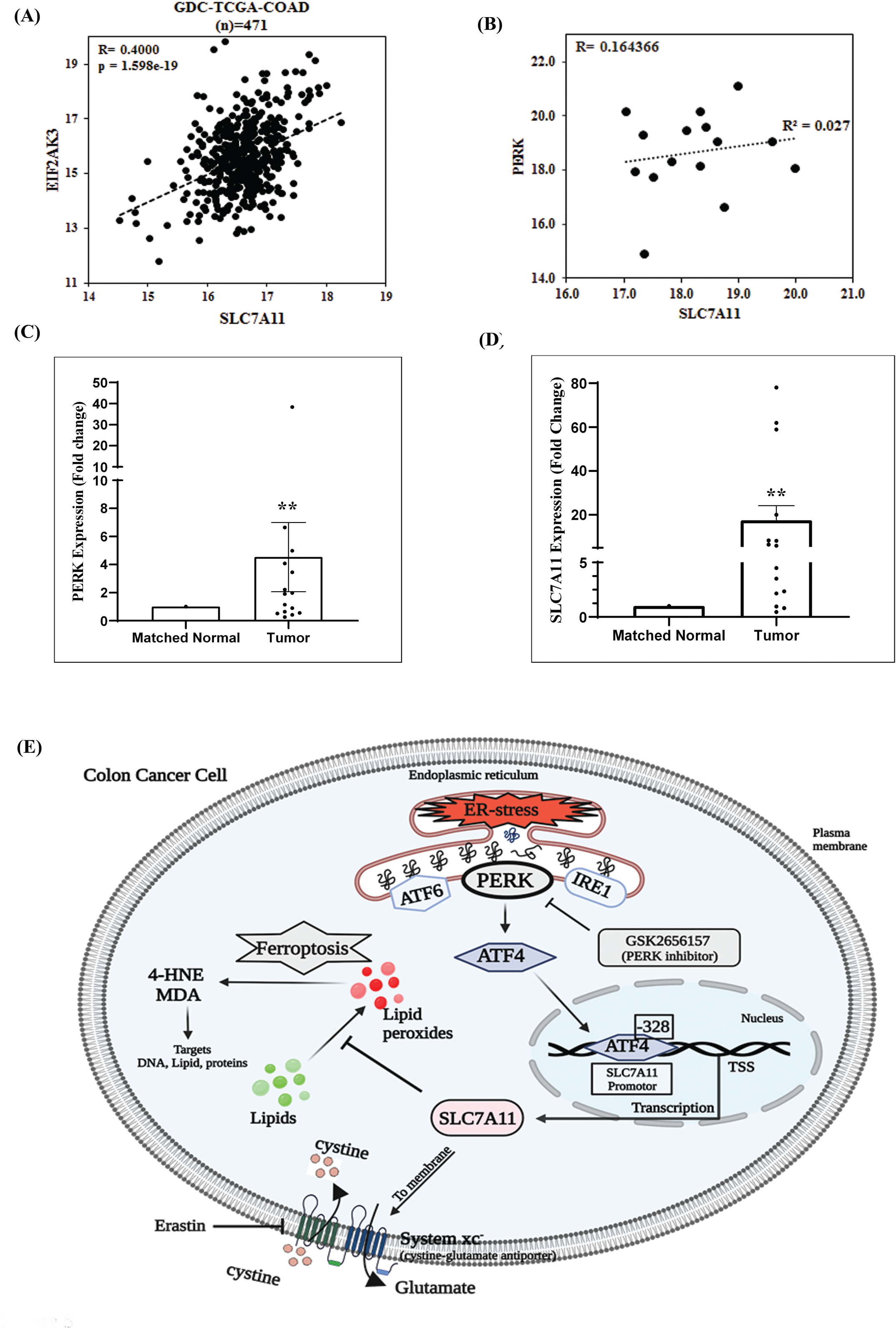
PERK (*EIF2AK3*) and *SLC7A11* are positively correlated in human colorectal tumors. (A) GDC TCGA COAD patient data were acquired from the Xena browser, and a correlation graph was plotted between *EIF2AK3* (PERK) and *SLC7A11*. R (Pearson’s correlation coefficient) (B) RT-PCR in matched non-malignant (Normal) and malignant tumor samples of colorectal cancer patients showed the correlation between delta Ct of *EIF2AK3* (PERK) and *SLC7A11.* (C-D) Total RNA was isolated from colorectal cancer patient tumor tissue samples along with their respective matched non-malignant counterparts, reverse transcribed, and RT-qPCR was performed for *PERK* and *SLC7A11* expression analysis. 18s is used as an internal control. Fold change in mRNA expression in (C) *PERK* and (D) *SLC7A11* is shown in bar diagram; Columns, the average value of fold change as compared to control; error bars ± SEM. *, p < 0.05, compared to control, n = 15. (E) The findings are illustrated in a graphical abstract showing how selectively PERK arm of ER-stress regulates ferroptosis in colorectal cancer.

The graphical abstract (Figure 5E) summarises how selectively the PERK arm of ER-stress/UPR protects cancer cells from ferroptosis. PERK mediated ATF4 upregulation and ATF4 binding to the promoter of *SLC7A11* results in upregulation of System Xc⁻ that stimulates cystine import into the cytosol and inhibits lipid peroxidation and ferroptosis.

## Discussion

Cancer cells make themselves resistant to therapeutic insults by smartly managing UPR and ER-stress responses. It has been well established in the literature how an apoptotic stimulus induces ER-stress in cancer cells leading to the activation of UPR and determine the final cell fate decisions [33–35]. However, these ER-stress and UPR responses are not well understood in the case of ferroptotic cell death. Our study highlights that ferroptosis inducers can cause UPR and ER-stress in colorectal cancer cells. However, PERK arm of UPR selectively plays a decisive role in modulating ferroptotic response [36,37]. In corollary with our findings, ferroptosis inducers like Sorafenib, Erastin, and 4-HNE (a by-product of lipid peroxides) have also been shown to induce ER-stress and UPR signaling [11,38–40]. Besides multiple reports demonstrating the involvement of the PERK arm of UPR in modulating apoptosis and autophagy, recent findings by Zheng et al. showed that PERK mediated sensitization of hepatocellular carcinoma in response to irradiation is due to enhanced apoptosis or ferroptosis [41]. In prostate cancer, it has been shown that loss of ATF6α promotes ferroptotic cell death, though the contribution of the other two arms (PERK and IRE1α) was not explored in this process [42]. In the current study, we observe the selective participation of PERK, over the other two UPR arms, in modulating ferroptosis. Our differential observations hint towards a possible contribution of context dependency and cancer specificity. Another unique aspect of our finding is that we observe exclusive ferroptotic cell death induction in the absence of PERK, which could only be rescued by ferroptosis inhibitor ferrostatin-1, and not by any other inhibitors of apoptosis and autophagy. This essentially advocates that the possibility of cross-talk between PERK mediated ferroptosis versus autophagy and apoptosis is weak when the upstream signals come from lipid peroxidation. In fact, any of the chemotherapeutics tested in our study failed to sensitize colorectal cancer to either apoptosis or ferroptosis in the absence of PERK. This further supports the exclusivity of PERK in modulating the ferroptotic process. In addition to this, PERK has been shown to be selectively activated during Epithelial-to-Mesenchymal Transition (EMT) process and making cancer cells vulnerable to ER-stress [43]. GPX4, SLC7A11 and ACSL4 are major downstream effectors of any ferroptotic process and the intricate balance between these play a critical role in executing full-blown ferroptosis [7,44,45]. For example, the presence of PERK in the cancer cells prevents the ferroptotic impact of GPX4 inhibitor RSL3 by positively regulating the expression of SLC7A11. Though the ferroptosis perspective was not studied, following exposure to paclitaxel, cancer cells have been shown to activate the PERK-ATF4 axis and maintain redox homeostasis by inducing the expression of the major antioxidant enzymes, including SLC7A11[46]. Several studies have indicated the regulation of *SLC7A11* by ATF4, but our study, for the first time, provided direct evidence for the transcriptional regulation of *SLC7A11* via ATF4 binding to its promoter [47,48]. In support of our observations, ATF4 has been shown to drive resistance to Sorafenib in hepatocellular carcinoma by preventing ferroptosis [49].

It is noteworthy to mention that in in vitro condition, functional loss of PERK either through PERK knockdown or by the treatment of PERK inhibitor alone does not result in ferroptotic cell death but loss of PERK function primes colorectal cancer cells towards ferroptosis which is evident by increased expression of 4-HNE and MDA in PERK knockdown state. However, in in vivo condition, we observe that PERK knockdown in colorectal cancer cells alone is able to reduce significant tumor growth and harvested tumors show robust positivity for ferroptosis markers like 4-HNE and MDA. The above results advocate that in in vivo condition under the influence of tumor microenvironment, PERK loss of function alone is sufficient to deliver its anti-tumor impact. Therefore, PERK inhibitors may have huge therapeutic potential in the clinics especially where tumors are non-responsive to chemotherapy that usually promotes apoptotic cell death.

Altogether, our study reveals a new role of the PERK-ATF4-SLC7A11 axis in modulating ferroptotic cell death in colorectal cancer in vitro and in vivo and posits therapeutic rationale for the development of small molecule PERK inhibitors against colorectal cancers that are commonly resistant to apoptosis but vulnerable to ferroptosis.

## Materials and Methods

### Reagents and antibodies

Dimethyl Sulfoxide (DMSO), ferrostatin-1, Doxorubicin, Paclitaxel, 5-fluorouracil, anti-β-Actin (cat#A3854) antibody, Doxycycline, Bovine Serum Albumin (BSA), and Polybrene were purchased from Sigma-Aldrich. RSL3 (cat#B6095) and Erastin (cat#B1524) were purchased from APExBIO. PERK inhibitor GSK2656157 (Cat#S7033) was purchased from Selleck Chemicals LLC. Recombinant Anti-Glutathione Peroxidase 4 antibody [EPNCIR144] (cat# ab125066) obtained from Abcam. Anti-ATF6α antibody clone# (37-1) (cat#73-505) was procured from BioAcademia. PVDF membrane and stripping buffer were obtained from Millipore Inc. BCA protein estimation kit, RIPA cell lysis buffer, blocking buffer, Super Signal West Pico and Femto chemiluminescent substrate, Lipofectamine-3000, Puromycin, FBS, RPMI-1640 media, Anti-Anti, were purchased from Thermo Fisher Scientific. Primers for real-time PCR and ChIP assay were purchased from IDT Inc and Eurofins Scientific. All chemicals were purchased from Sigma or Thermo scientific unless specified otherwise. Antibodies were obtained from cell signaling technology (CST) or mentioned otherwise.

### Cell Culture

Colorectal cancer cell lines HT29 and SW620 were obtained from American Type Culture Collection (ATCC), USA. Mycoplasma-free early passage cells were resuscitated from liquid nitrogen vapor stocks and inspected microscopically for stable phenotype before use. HT29 cells were cultured in RPMI-1640 medium containing 10% fetal bovine serum (Gibco/Invitrogen), supplemented with anti-anti (Invitrogen containing 100 μg/ml streptomycin, 100 unit/ml penicillin, and 0.25 μg/ml amphotericin B). SW620 cells were cultured in DMEM medium containing 10% fetal bovine serum (Gibco/Invitrogen), supplemented with anti-anti. RPMI-1640 L-cysteine, L-cystine, L-glutamine and L-methionine free media (MP Biomedicals) was used to culture HT29 cells in cystine-free conditions. STR profiling was performed to authenticate all the cell lines employed in the investigation. Cell lines were cultured in an Eppendorf Galaxy 170R/170S CO2 incubator to provide a stable and homogeneous 5% CO_2_ and 37°c temperature and humid atmosphere required for cell culture.

### Cytotoxicity assay (SRB assay)

In-vitro cytotoxic effects of RSL3, Paclitaxel, Doxorubicin, 5-Fluorouracil, and Oxaliplatin were assessed with standard SRB (Sulforhodamine B) assay as described before [50,51]. Following an incubation period of 48 hours, cell monolayers were fixed with 10% (wt/vol) trichloroacetic acid and stained for 30 minutes before being washed repeatedly with 1% (vol/vol) acetic acid to remove excess color. The protein-bound dye was dissolved in a 10 mM Tris base solution, and the absorbance of the treated and untreated cells was measured on a multi-well scanning spectrophotometer (Epoch Microplate Reader, Biotek, USA) at a wavelength of 510 nm. All the calculations for percent inhibition and IC50 were done in excel.

### Western blotting

After harvesting, the cells or tissues were lysed on ice with Pierce™ RIPA lysis solution for 30 minutes. The Pierce™ BCA protein assay kit was used to estimate the protein concentration of the lysates. Equal quantity of the protein was resolved in a 4–15% Mini-PROTEAN® TGX™ Precast Protein Gels and transferred to an Immun-Blot PVDF Membrane (Bio-Rad) for antibody incubation. After transfer, the PVDF membrane was blocked with 5% non-fat dry milk or 5% BSA, followed by incubation with appropriate dilutions (manufacturer’s protocol) of primary antibodies overnight at 4°C. After 3 washes for 5 minutes each, the membrane was incubated with a 1:5000 dilution of horseradish peroxidase-conjugated secondary antibody for hour at room temperature. Immunoreactivity was detected by enhanced chemiluminescence solution (Bio-Rad Clarity Western ECL Substrate and Immobilon^TM^ western, Millipore, USA) and scanned by the gel documentation system (Bio-Rad chemidoc XRS plus).

### Flow cytometry

We used flow cytometry to assess the cytotoxic effect of RSL3, Paclitaxel, Doxorubicin, 5-Fluorouracil, Oxaliplatin in HT29 EV (untagged) and PERK knockdown cells (pUltra-Chili-Luc-red). In brief, 6 lacs cells/well were seeded in 6-well plates and allowed to grow with treatment or vehicle control. Cells were harvested with TrypLE (Invitrogen) for single-cell suspension in FACS buffer (PBS with 0.1% BSA and 1mM EDTA). After washing and centrifugation, cell pellets were resuspended in FACS buffer and analysed by FACS Calibur (BD). Acquired data were analysed using FlowJo software (Treestar).

### BODIPY 581/591 C11 Assay

BODIPY 581/591 C11 (Invitrogen) is a lipid-soluble fluorescent indicator of lipid peroxidation. Upon oxidation, its excitation maximum shifts from 581 to 500 nm, and the emission maximum shifts from 591 to 510 nm. To estimate lipid peroxidation in vitro, 6 lacs cells per well were seeded in 6-well plates and treated with RSL3 for 6 hours or mentioned otherwise. After completion of treatment, cells were incubated with 5μM BODIPY C11 in the CO_2_ incubator for 30 minutes. After incubation, cells were harvested with TrypLE, washed with DPBS, and analysed by FACS Calibur (BD). Acquired data were analysed using FlowJo software.

### Generation of stable cell lines by lentiviral transduction

3rd generation lentiviral vector pUltra-Chili-Luc (addgene no. 48688) with the bi-cistronic expression of tdTomato and luciferase was used to make HT29 cells fluorescent tagged. Lentiviral particles were generated in HEK-293T cells. Transduction was carried out in the presence of Polybrene (8 μg/ml). A population of transduced cells (HT29-Chili-Luc) was identified by chilli red expression and sorted by flow cytometry. For *PERK, SLC7A11*, and *ATF4* knockdown generation, shRNA sequences were cloned into the 3rd generation transfer plasmid pLKO.1 TRC cloning vector (Addgene cat no. 10878) between unique AgeI and EcoRI sites downstream of the U6 promoter. HEK-293T cell line was used to generate lentiviral particles using the transfection reagent Lipofectamine 3000. The media containing the viral particles was supplemented with Polybrene (8 μg/ml) for transduction. Cells were subjected to puromycin selection after 48 hours of transduction, and the knockdown profile of PERK, SLC7A11, and ATF4 was confirmed after 1 week of selection via western blot. Following oligo sequences were used to clone *PERK, SLC7A11* and *ATF4* shRNA in pLKO.1 plasmid-

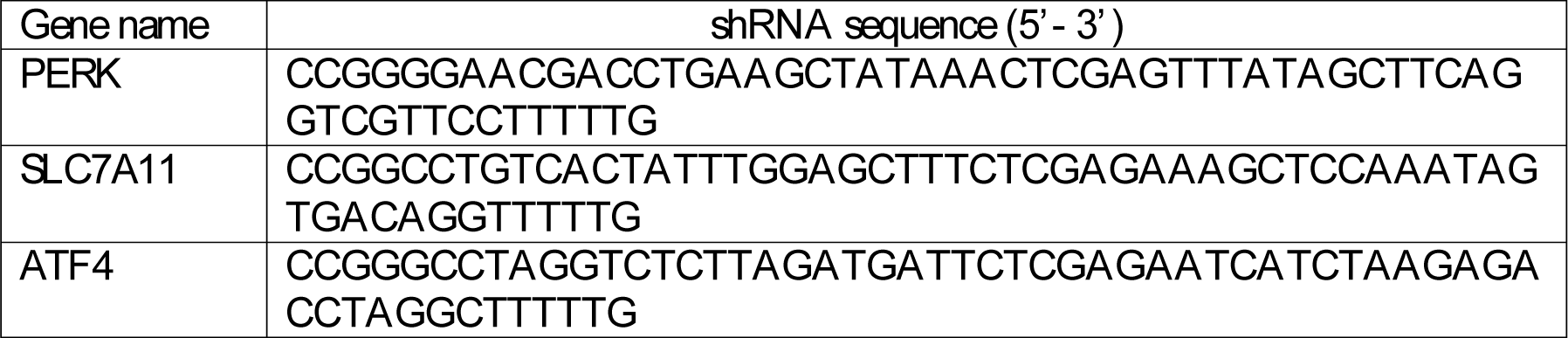

### Real-time quantitative PCR (RT-qPCR)

Total RNA was isolated from the cultured cells and tissues using the standard procedure of the RNeasy Mini Kit (Qiagen, cat no.74104). The concentration and purity of the RNA samples were determined using nanodrop. Total RNA (1μg) of each sample was reverse-transcribed (RT) with random hexamer according to the manufacturer’s protocol (Verso cDNA synthesis kit). The final cDNA was diluted with nuclease-free water (1:5), and 10ng of diluted cDNA was used for each reaction in real-time PCR. Real-time PCR was carried out using an ABI 7500 Real-Time PCR System (Applied Biosystems). Reactions for each sample were performed in triplicate. 18s amplification was used as the housekeeping gene control. The Standard delta-delta Ct method was used to calculate the relative fold change in gene expression. For amplifying *PERK*, *ATF4*, and *SLC7A11*, we performed SYBR Green-based RT-PCR following the manufacturer’s (PowerUp SYBR Green ABI) instructions.

### Chromatin Immunoprecipitation (ChIP) Assay

Following the manufacturer’s protocol, ChIP assay was conducted using the CUT&RUN Assay Kit (cat# 86652 Cell Signaling Technology). In brief, cells at 80% confluency were fixed with 1% formaldehyde for 10 minutes. Cells were then centrifuge washed, followed by lysis in 200μl of membrane extraction buffer containing protease inhibitor cocktail. The cell lysates were digested with MNase for 30 minutes at 37°C to get chromatin fragments of 100-500 bp long DNA fragments. Incubation with ATF4 rabbit monoclonal primary antibody, normal IgG, and anti-histone 3 (H3) was done overnight at 4°C. After washing with the wash buffer three times, elution of chromatin from Antibody/Protein A Magnetic beads and reversal of cross-linking was carried out by heat. Spin columns purified DNA then used for SYBR Green-based real-time PCR. The following primers were used to amplify the −328bp site on *SLC7A11* promoter forward primer- 5’CTACTCACAAAACAGTCGCA3’, reverse primer- 5’GCAACTCGTAGTGAGCAACAA3’, and −494bp and −447bp sites were amplified using forward primer- 5’ATTGGATTTGACTGTATTGCCTT3’ and reverse primer- 5’CATTGTTTATAACAACACAGTTTGA3’.

### Cloning of *SLC7A11* promoter in luciferase reporter vector and luciferase assay

The following primers were used to PCR amplify the *SLC7A11* promoter −0.6 kb upstream of the TSS from the HEK293 cell line-forward primer- 5’TCGGCTAGCGAGGAAGGCTTATAGTTGTGTGTATGTGAC3’, reverse primer-5’AGCCTCGAGCAGCTCAGCTTCCTCATGGGC3’. The amplified fragments were cloned into the PGL4.12 [luc2 CP] vector between the NheI and XhoI restriction sites. HT29 cells were seeded in a 6-well plate upto 50-60% confluency and transfected with 2.5μg of cloned PGL4.12 and 50ng of PGL4 (hRluc-CMV) plasmid using lipofectamine-3000 as transfection reagent (Invitrogen). The transfected cells with cloned pGL4.12 and pGL4 (hRluc-CMV) were treated with vehicle control or two doses (5μM and 1μM) of PERK inhibitor (GSK2656157) for 24 hours. Cells were lysed with 100μL of lysis buffer provided with the Dual-Glo Luciferase assay kit (Promega). The GloMax® 96 Microplate Luminometer was used to measure the activity of Firefly and Renilla luciferases according to the manufacturer’s protocol (Promega). For each sample, firefly luciferase activity was normalised to Renilla luciferase activity, and fold change in luciferase activity in different treatment groups was calculated.

### In vivo studies in xenograft tumor models

All animal studies were conducted following standard principles and procedures approved by the Institutional Animal Ethics Committee (IAEC) of CSIR-Central Drug Research Institute. All the animals were maintained in a pathogen-free facility under a day-night cycle. Mice were randomly assigned to groups by a blinded independent investigator. Following our well-established colorectal cancer xenograft models[3], we inoculated 2 × 10^6^ cells (HT29 EV and HT29 PERK KD each) in 100μl PBS subcutaneously into the flanks of the left or right hind leg of each 4–6 weeks old nude Crl: CD1-Foxn1^nu^ mice. Throughout the study, the tumors were measured with an electronic vernier caliper at regular intervals, and the tumor volumes were calculated using the standard formula V = (W (2) × L)/2, where ‘W’ is the short and ‘L’ is the long tumor axis. At the end of the experiment, mice were sacrificed, and subcutaneous tumors were dissected for further studies.

### Immunohistochemistry of tumor tissues

Harvested tumors were fixed in 10% neutral buffered formalin (NBF), and paraffin blocks were prepared for sectioning. For staining, tissue sections were deparaffinised and rehydrated by passing through serial dilutions of xylene and ethanol. Antigen retrieval was performed in 10mM sodium citrate buffer (pH 6.2) by heating at 95°C for 20-30 minutes. Processed slides were washed in PBS for 5 minutes. We used VECTASTAIN ABC KIT (VECTOR laboratories) for IHC staining. ImmEdge pen (hydrophobic barrier pen), and Bloxall blocking solution, were purchased from Vector Laboratories, Inc. Fluorochrome conjugated secondary antibodies, ProLong™ Gold Antifade Mountant, were purchased from molecular Probes-Invitrogen. The endogenous peroxide activity was neutralised by incubating the slides with BLOXALL blocking solution for 10 minutes. After washing, tissue sections were incubated with diluted normal blocking buffer for 20 minutes to prevent non-specific staining. The sections were incubated overnight at 4°C with primary antibodies of 4-HNE and MDA diluted in buffer (1:100). The sections were incubated with biotinylated secondary antibody for 1 hour, followed by VECTASTAIN ABC Reagent for 30 minutes. Slides were incubated with diaminobenzidine (DAB) as a chromogen and counterstained with hematoxylin. Negative control sections were processed in parallel without the primary antibody incubation. The sections were dehydrated and mounted using DPX (Sigma). Stained sections were examined under a microscope (EVOS XL core) under 10x and 40x magnification. ImageJ software was used for scoring 4-HNE and MDA staining. The color deconvolution tool split the IHC images into three color images separately, showing different staining intensities. After adjusting the threshold, images were changed in binary. Analyse the particle tool was used to get the average value of pixels in the DAB channel. ATM score was calculated following the standard formula (Refs) ATM Score = 1/255 (the average value of all the pixels in the DAB channel), where ATM stands for Average Threshold Measure, and 255 is the value of maximum staining intensity [52,53].

### Analysis of TCGA colorectal cancer dataset

Illumina HiSeq mRNA data from GDC TCGA colorectal Cancer (n=471) was downloaded from the TCGA portal for *PERK* and *SLC7A11* genes using the UCSC Xena browser (https://xena.ucsc.edu) [54]. Log2 (fpkm-uq + 1) values for *PERK* and *SLC7A11* were used to draw a scatter plot. Pearson’s correlation coefficient (r) and p-value were calculated in excel.

### Patient samples collection and RNA isolation

Patient samples were scrutinised following the set criteria. A total of 15 CRC tumors with paired normal colorectal mucosa samples were collected from RGCIRC (Rajiv Gandhi Cancer Institute and Research Centre), India, from 2016 to 2019. It was ensured that the tumor was sporadic and the patient had not received chemotherapy or radiotherapy before surgery. The pathologist ensured the total oncogenic area of cancerous cells was not less than 80%. Informed consent was obtained from each patient. The ethical approval of the study was approved by the Institute Ethics Committee, Motilal Nehru National Institute of Technology Allahabad (Ref. No. IEC17-18/027). Total RNA was extracted from tumor tissues using RNeasy Mini Kit (Qiagen, cat no.74104), and RNA concentration and quality were estimated using nanodrop (BioTek Take3). The cDNA preparation and RT-PCR analysis were done as described above.

### Statistics

In the figure legends, most in vitro experiments represent at least two or more independent experiments or specified otherwise. Student’s t-test was used to examine statistically significant differences for two-group analysis. All data are presented as means ± SD or SEM. These analyses were done with Graph-Pad Prism software. Results were considered statistically significant when p values ≤ 0.05 between groups.

## Availability of Data and Materials

All data needed to evaluate the conclusions in the paper are present in the paper and/or in the Supplementary Information. Additional data related to this manuscript will be made available upon reasonable request.

## Acknowledgments

We sincerely acknowledge the excellent technical help of Mr. A. L. Vishwakarma of SAIF for the Flow Cytometry studies; Authors are immensely grateful to Dr. Juhi Tayal, Dr. Anurag Mehta, Dr. DC Doval and Ms. Somika Tiwari (Biorepository, Rajiv Gandhi Cancer Institute and Research Centre, New Delhi, India) for colorectal tumor samples. Research of all the authors’ laboratories was supported by CSIR Pan-Cancer Grant (HCP-40) and Fellowship grants from CSIR, DBT, and UGC. D.D. acknowledges grant support from CSIR-FTT (MLP-2025), DST (EMR/2016/006935), DBT (BT/AIR0568/PACE-15/18) and ICMR (2019-1350). The Institutional (CSIR-CDRI) communication number for this article is 135.

## Author Contributions

KKS was involved in study designing, performed experiments and wrote the draft manuscript. MPS performed bioinformatic analysis. PC, ABS, MAK AV, MAN, SRS, SM, and AS helped in carrying out in vitro and in vivo studies. SS provided support for patient sample data generation. JS helped in animal maintenance and experimentation. DD conceived the idea, designed experiments, analysed data, wrote the manuscript and provided overall supervision. All authors read and approved the final manuscript.

