## Supplementary Figure for "PERK arm of UPR selectively regulates ferroptosis in colon cancer cells by modulating the expression of SLC7A11 (System Xc-)"

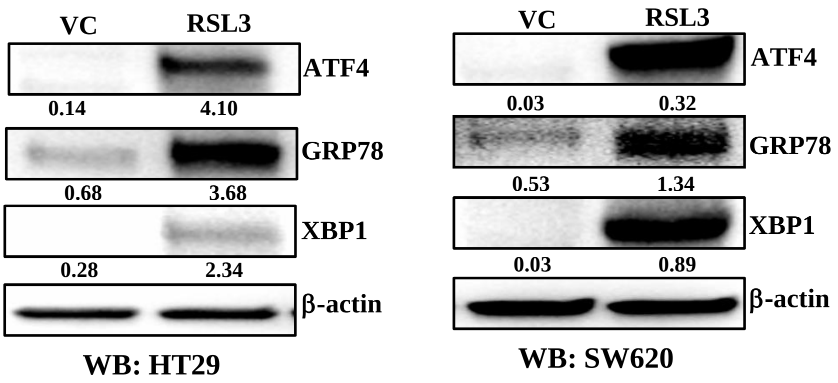


**Supplementary Figure 1. RSL3 promotes UPR downstream responses in colon cancer.**

HT29 (left) and SW620 (right) cells were either treated with 1μM RSL3 or vehicle control (VC) for 24 hours and protein lysates were prepared for western blot analysis. Immunoblot shows the expression for UPR downstream effector proteins like ATF4, Bip/GRP78 and XBP1.

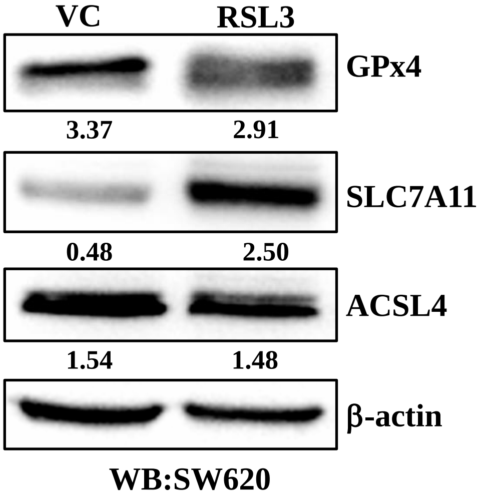


**Supplementary Figure 2. RSL3 promotes SLC7A11 expression in colon cancer.**

SW620 cells were either treated with 1μM RSL3 or vehicle control (VC) for 24 hours and protein lysates were prepared for western blot analysis. Immunoblot shows the expression for major ferroptosis regulator proteins GPx4, SLC7A11, and ACSL4 in SW620 cells.

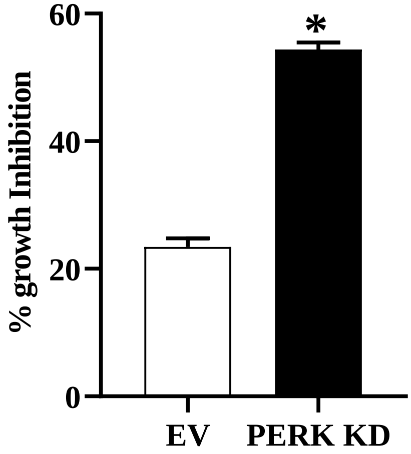


**Supplementary Figure 3. PERK loss of function sensitizes Erastin response in colon cancer.**

Control (EV) and PERK knockdown HT29 cells were treated with vehicle or 12.5µM erastin for 48 hours and the SRB assay was performed. Percent growth inhibition was tabulated. *$p < 0.05$; compared to EV.

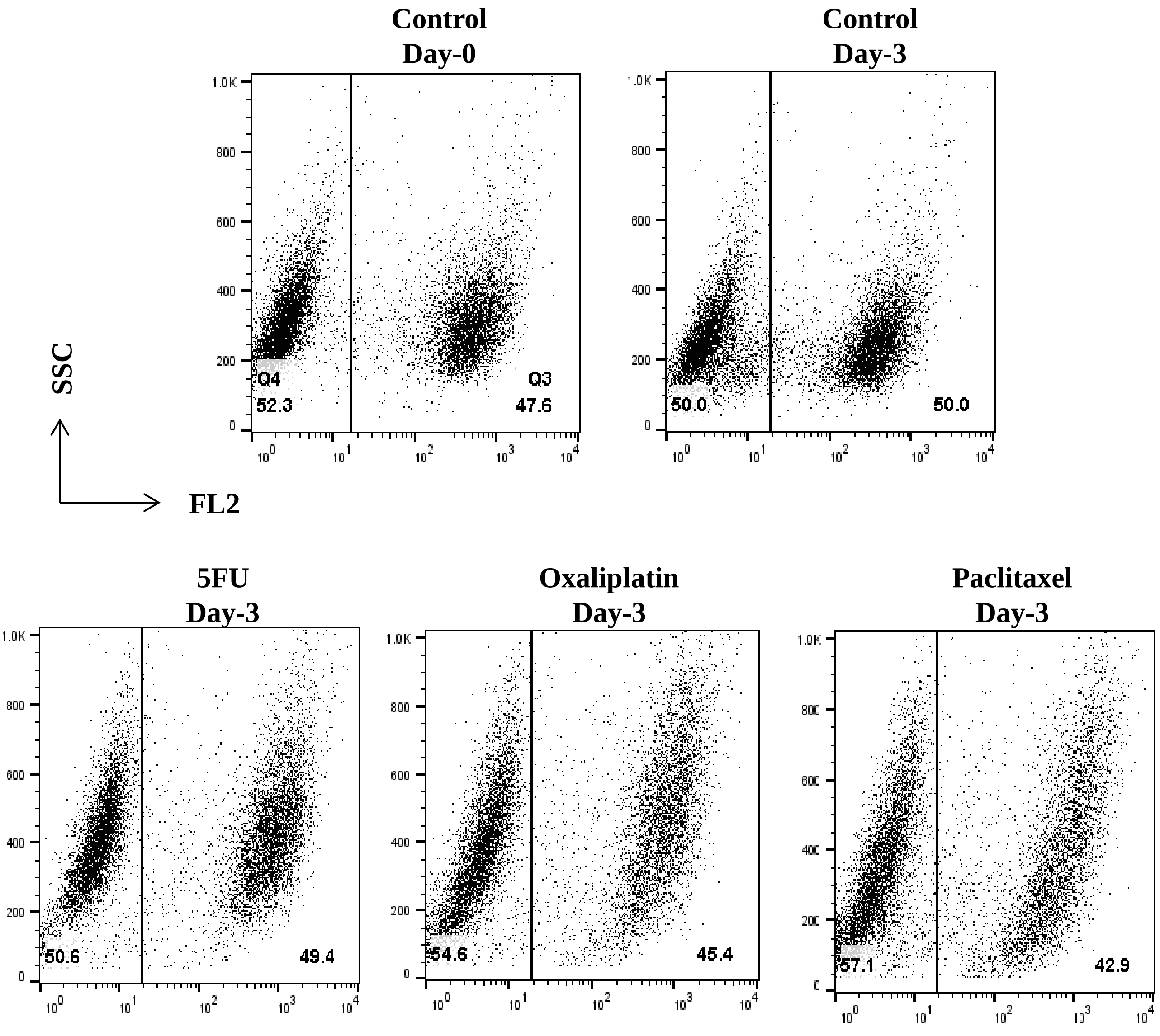


**Supplementary Figure 4. PERK loss of function does not sensitize chemotherapy response in colon cancer.**
HT29 EV (untagged) and chilli-tagged PERK KD (red) cells were mixed in equal numbers and subjected to flow cytometric analysis on day 0 and day 3 following treatment with vehicle control or 5-Fluoruracil (100µM), Oxaliplatin (10µM) or paclitaxel (10nM). chilli-tagged PERK KD cells are gated in the right quadrant and % positivity are noted at the bottom.

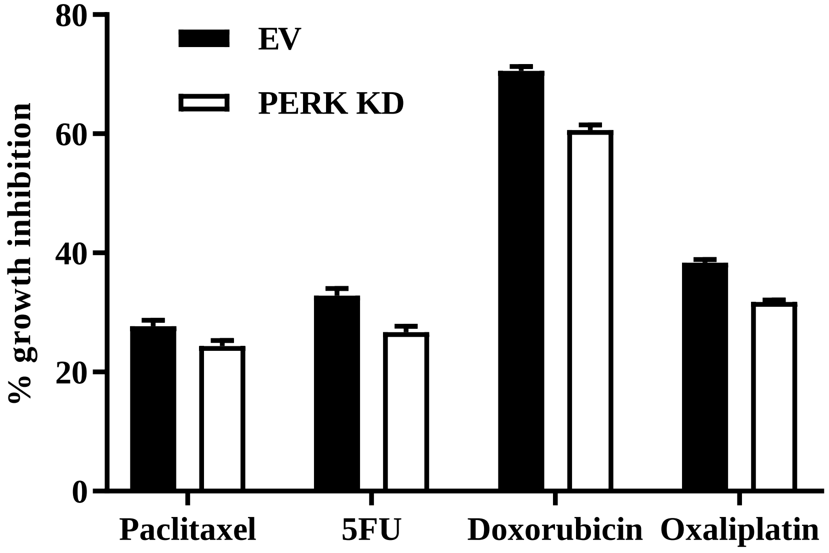


**Supplementary Figure 5. PERK loss of function does not sensitize chemotherapy induced cytotoxic response in colon cancer.**

Control (EV) and PERK KD SW620 cells were treated with either vehicle control or with different chemotherapeutic drugs paclitaxel (20nM), 5-Fluorouracil (100μM), Doxorubicin (10μM), Oxaliplatin (50μM) for 48 hours and cytotoxic impact of these drugs was evaluated via SRB assay. Percent growth inhibition was tabulated.
